# The Synergism Between DHODH Inhibitors and Dipyridamole Leads to Metabolic Lethality in Acute Myeloid Leukemia

**DOI:** 10.1101/2020.10.10.334631

**Authors:** Valentina Gaidano, Mohammad Houshmand, Nicoletta Vitale, Giovanna Carrà, Alessandro Morotti, Valerio Tenace, Stefania Rapelli, Stefano Sainas, Agnese Chiara Pippione, Marta Giorgis, Donatella Boschi, Marco Lucio Lolli, Alessandro Cignetti, Giuseppe Saglio, Paola Circosta

**Author notes:** **Authors’ information**, Mohammad Houshmand and Nicoletta Vitale contributed equally to this work. Giuseppe Saglio and Paola Circosta contributed equally to this work.

## Abstract

**Background:** Dihydroorotate Dehydrogenase (DHODH) is a key enzyme of the *de novo* pyrimidine biosynthesis, whose inhibition was recently found to induce differentiation and apoptosis in acute myeloid leukemia (AML). DHODH inhibitors were previously investigated in solid tumors, where they showed promising antiproliferative activity, both *in vitro* and *in vivo*. However, their effectiveness was not confirmed in clinical trials, probably due to the pyrimidine salvage pathway that cancer cells could exploit to survive. In this study we investigated the pro-apoptotic activity of MEDS433, the DHODH inhibitor developed by our group, against AML. Learning from previous failures, we challenged our model mimicking *in vivo* conditions, and looked for synergic combination to boost apoptosis.

**Methods:** We evaluated the apoptotic rate of multiple AML cell lines and AML primary cells treated with MEDS433 or other DHODH inhibitors, alone and in combination with classical antileukemic drugs or with dipyridamole, a blocker of the pyrimidine salvage pathway. Experiments were also performed mimicking *in vivo* conditions, i.e., in the presence of physiological uridine plasma levels (5 *μ*M).

**Results:** MEDS433 showed a strong apoptotic effect against multiple AML cell lines, which was at least partially independent from the differentiation process. Its combination with classical antileukemic agents resulted in a further increase of the apoptotic rate. However, when MEDS433 was tested in the presence of 5 *μ*M uridine and/or in primary AML cells, results were less impressive. On the contrary, the combination of MEDS433 with dipyridamole resulted in an outstanding synergistic effect, with a dramatic increase of the apoptotic rate both in AML cell lines and AML primary cells, which was unaffected by physiological uridine concentrations. Preliminary analyses show that the toxicity of this treatment should be limited to proliferating cells.

**Conclusions:** The combination of a DHODH inhibitor and dipyridamole is characterized by differentiating and pro-apoptotic features and induces metabolic lethality on a wide variety of AMLs with different genetic backgrounds.

## 1 Introduction

The inhibition of Dihydroorotate Dehydrogenase (DHODH) has recently been found to induce differentiation in several models of Acute Myeloid Leukemia (AML), both *in vitro* and *in vivo* [1]. From this seminal discovery, several academic and industrial research groups, including ours, have designed new and more potent DHODH inhibitors, confirming original results and extending the knowledge about this topic [2, 3, 4, 5].

Transcriptome analyses revealed that leukemic cells treated with DHODH inhibitors upregulate genes related to apoptosis and differentiation, and downregulate protein translation-related genes, impairing protein synthesis [4, 5]. However, the exact mechanisms triggered by DHODH inhibitors that eventually affect differentiation and apoptosis have not been fully elucidated [6], probably because DHODH is involved in multiple cellular pathways, and its inhibition has very wide consequences. In particular, DHODH is a mitochondrial enzyme which catalyses the conversion of dihydroorotate to orotate; it is strictly associated with the electron transport chain and, simultaneously, a fundamental enzyme in the *de novo* pyrimidine biosynthesis [7]. Evidences [8] suggest that DHODH inhibition acts through pyrimidine starvation, rather than cellular respiration impairment; the counterproof is that uridine, a downstream product of DHODH in the pyrimidine biosynthesis, is able to abolish the effect of DHODH inhibitors on AML cell lines [1].

Unlike differentiation, DHODH inhibition-induced apoptosis has not been thoroughly investigated per se; it has probably been considered a consequence of differentiation, somehow similarly to the effect induced by all-trans retinoic acid (ATRA) in promyelocytic leukemia. However, DHODH inhibitors were initially tested in solid tumors, where they showed to reduce the tumor growth *in vitro;* DHODH inhibitors were also found to increase p53 levels, and to synergize with an inhibitor of p53 degradation to reduce tumor growth *in vivo* [9]. Indeed, pyrimidines are crucial for the proliferation of living entities, representing the basic constitutive elements for DNA and RNA, and being involved in the synthesis of glycoprotein, glycolipid and phospholipid synthesis [6]. Accordingly, the depletion of intracellular pyrimidine pool results in cell cycle arrest in S-phase [9].

In spite of the underlying mechanisms, DHODH inhibition is an extremely interesting strategy, as *i*) it represents a new pathway to fight AML; ii) differentiation induction has yielded incredible results in acute promyelocytic leukemia, being potentially able to induce differentiation of leukemic stem cells, thus reducing the relapse rate; iii) preliminary studies do not show a significant toxicity, potentially allowing DHODH inhibitors to be combined with other drugs. Not surprisingly, 4 different agents are currently being tested in clinical trials for AML, including old (brequinar) and new compounds (PTC299, ASLAN003, BAY2402234).

Despite this justified enthusiasm, there are also few concerns about the efficacy of DHODH inhibitors, especially if they are used alone.

DHODH inhibitors have already failed to prove their efficacy in clinical trials on solid tumors [10, 11, 12]. One of the major causes of their failure directly derives from their mechanism of action, i.e., the pyrimidine starvation. There are two pathways that guarantee the required pyrimidines to the cells: the *de novo* pathway, where DHODH plays a crucial role, and the salvage mechanism, where uridine and cytidine are acquired from the extracellular space and recycled. Resting cells generally obtain pyrimidines with the salvage mechanism, while rapidly growing cells have to rely on the *de novo* pyrimidine biosynthesis, in order to get large amounts of nucleotides [13]. This is probably the reason why DHODH inhibitors have a low toxicity profile in non-proliferating cells; the other side of the coin is that the salvage pathway basically represents a natural escape from DHODH inhibitors that cancer cells could leverage to survive: uridine, in fact, is naturally present in plasma and can enter the cells through hENT1/2 (Equilibrative Nucleoside Transporters) or, to a lesser extent, hCNT (Concentrative Nucleoside Transporters) channels. Accordingly, *in vivo* experiments with DHODH inhibitors on AML generally show less impressive results compared to *in vitro* settings [1]. Further concerns derive from the notion that AML is an extremely aggressive cancer, whose biology is characterized by the subsequent emergence of resistant subclones; for this reason, AML is generally treated with multidrug therapies, whether chemotherapy-based (polychemotherapy) or non-chemotherapy based (e.g. hypomethylating agents + bcl2 inhibitors).

In order to improve DHODH inhibitors performances and look for synergic combinations, we decided to test *in vitro* multiple drug combinations, starting with the most commonly used drugs in AML. Moreover, we hypothesized that the contemporary blockade of both the *de novo* and the salvage pyrimidine pathways could be fatal to cancer cells, which are dependent on pyrimidine supply. In particular, we associated MEDS433, the DHODH inhibitor developed by our group, to dipypiridamole, an antiplatelet drug with vasodilating properties which is also known to inhibit hENT1 and hENT2 channels, thereby greatly reducing the nucleoside/nucleotide influx of the salvage pathway.

In summary, this work aims to: i) characterize DHODH inhibition-induced apoptosis, analyzing the effect of MEDS433 on different AML cells and in different conditions, and ii) find synergic combinations to boost the activity of DHODH inhibitors.

## 2 Results

### 2.1 MEDS433 induces apoptosis in several AML cell lines

MEDS433 is a new, potent DHODH inhibitor, developed and characterized by our group, which can induce differentiation in multiple AML cell lines, at a 1-log lower concentration compared to brequinar [2]. Here we show that MEDS433 also has a strong apoptotic effect on several AML cell lines, again at a 1-log lower concentration compared to brequinar (Fig. 1A). In particular, MEDS433 was effective on THP1, U937, NB4 and, to a lesser extent, on OCI AML3 and MV4-11 cell lines, where it induced apoptosis (Fig. 1) and a dramatic drop in the number of viable cells (Fig. S1A, Fig. S1B). The apoptotic effect was totally abrogated when uridine was added to the culture at hyperphysiological concentrations (Fig. 1B), indicating that apoptosis was indeed due to pyrimidine starvation rather than off-target effects. MEDS433 pro-apoptotic and pro-differentiating activity was conserved in niche-like conditions, i.e., when AML cell lines were co-cultured with a stromal cell line or in hypoxic conditions (Fig. 1D and Fig. S1C). Tab. S1 shows the molecular characterization of utilized AML cell lines.

**Figure 1:**
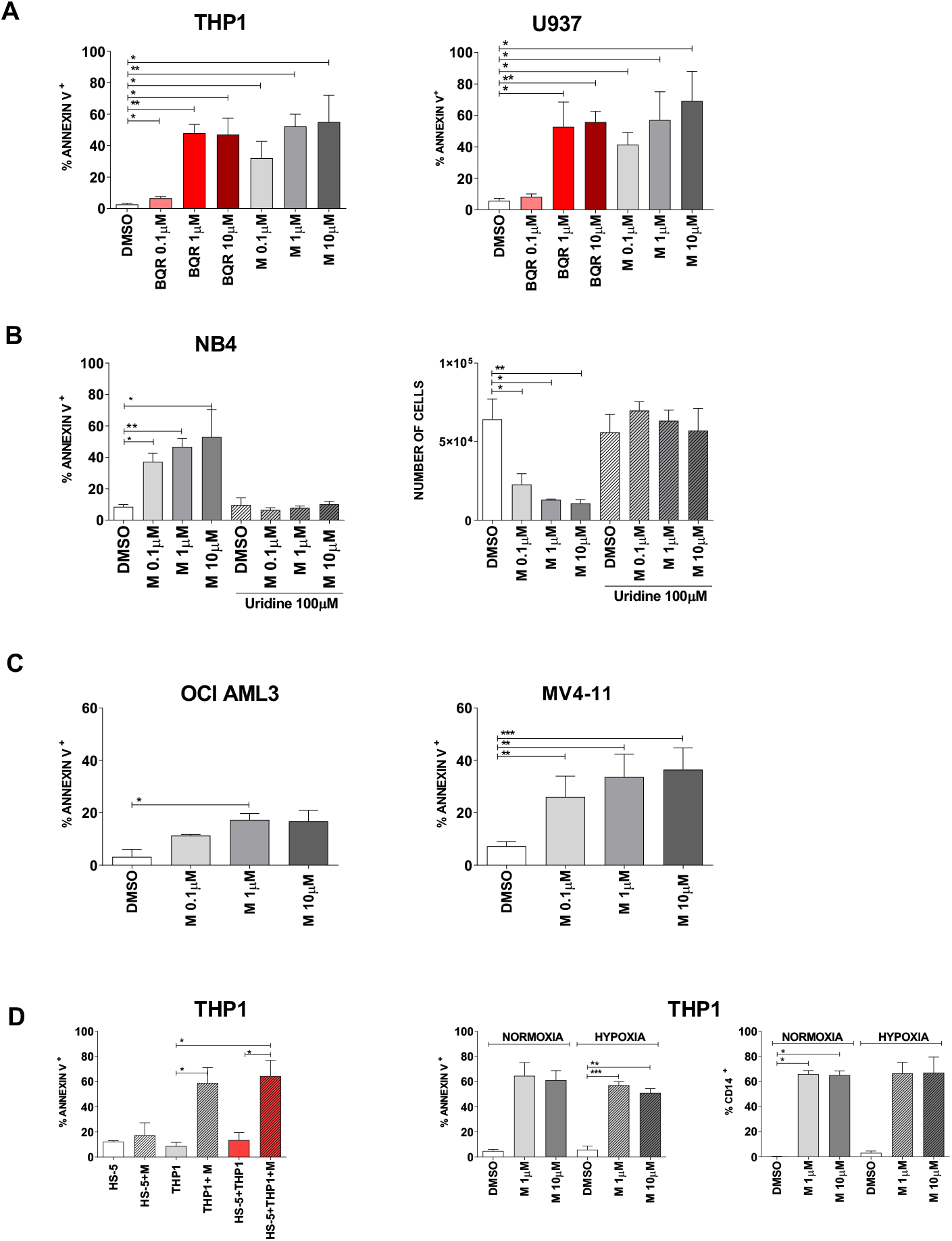
Apoptosis induced by various DHODH inhibitors in different AML cell lines and in different conditions. (A) Comparison between the apoptotic rate induced by MEDS433 and brequinar at different concentrations. (B) Hyperphysiological levels of uridine (100 *μ*M) abrogate the apoptotic effect (left panel), preserving the number of viable cells (right panel). (C) OCI AML3 and MV4-11 have a reduced sensitivity to MEDS433 compared to THP1, U937 and NB4 cell lines. (D) MEDS433 activity is preserved in niche-like conditions: the left panel shows the apoptotic rate induced by MEDS433 at 1 *μ*M on THP1 cells alone or co-cultured with HS-5, a stromal cell line; the middle and right panel show, respectively, the apoptotic rate and the differentiation effect induced by MEDS433 in normoxic and hypoxic conditions. In all experiments, apoptosis and differentiation were evaluated after 3 days of treatment. Apoptosis was evaluated through Annexin V expression; cell differentiation was evaluated through CD14 expression. DMSO indicates cells treated with dimethyl sulfoxide only. M: MEDS433. BQR: brequinar. Ur: uridine. Statistical significance: t-test, *p < 0.05; **p < 0.01; ***p < 0.001.

### 2.2 Apoptosis is time-dependent, allowing to rescue partially resistant cell lines

Different cell lines show different sensibility to DHODH inhibition (Fig. 1); we hypothesized that cells with low proliferating rates could be less sensitive to DHODH inhibitors, as they could still rely on the salvage pathway for pyrimidine supply. Accordingly, Fig. 2A suggests that the apoptotic rate correlates with cell lines’ doubling time (r2=0.3). However, when partially resistant cells were treated for 6 instead of 3 days, apoptosis was significantly increased and the number of viable cells was remarkably reduced compared to control (Fig. 2B and Fig. 2C).

**Figure 2:**
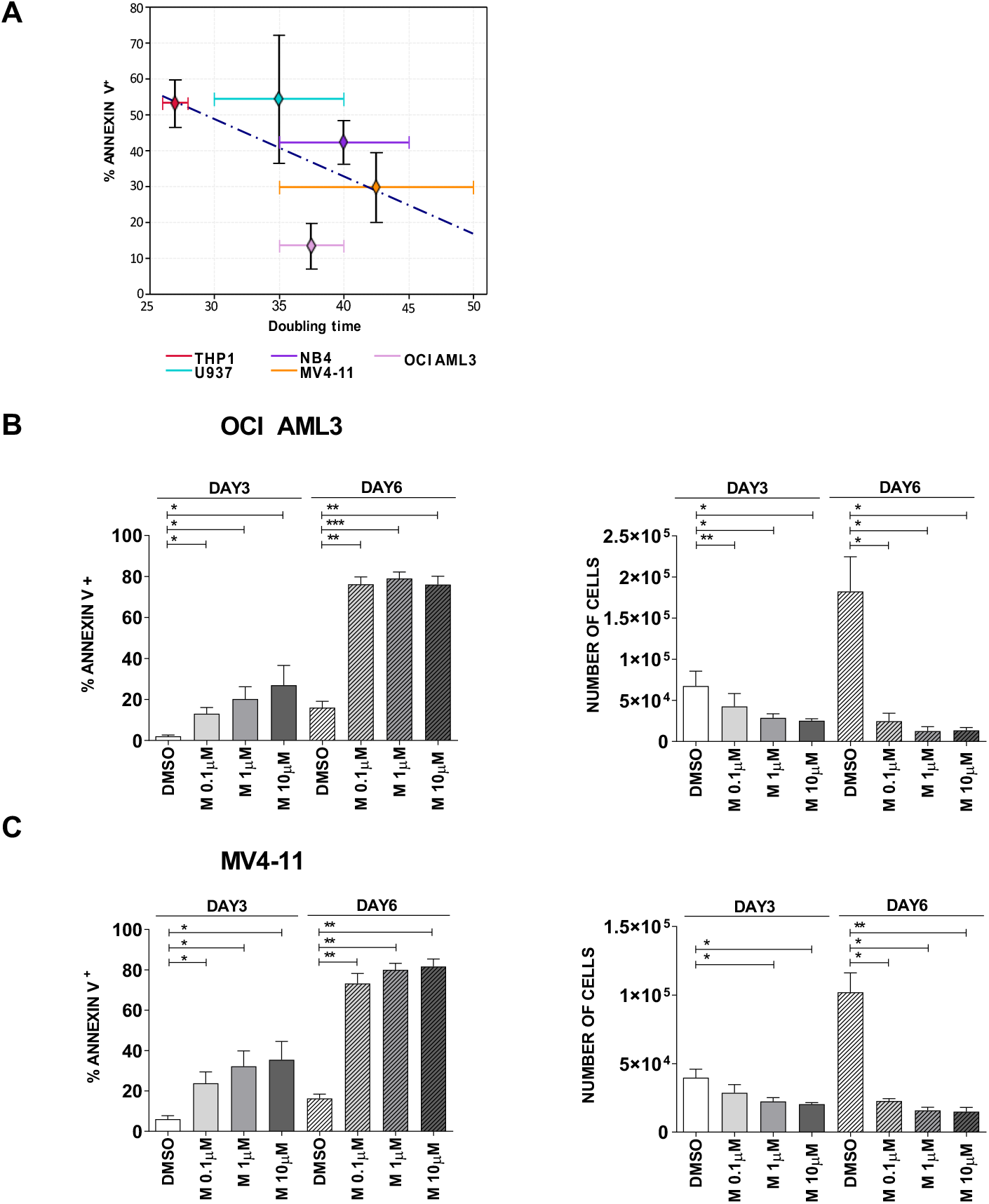
Rescue of less sensitive cell lines. (A) Regression line correlating the apoptotic rate induced by MEDS433 at 3 days and the doubling time of tested AML cell lines. (B and C) Apoptotic rate and corresponding viable cell counts of OCI AML3 (B) and MV4-11 (C) treated for 3 or 6 days with MEDS433 at different concentrations. DMSO: dimethyl sulfoxide. M: MEDS433. Statistical significance: t-test, *p < 0.05; **p < 0.01; ***p < 0.001.

### 2.3 Apoptosis is both the result of differentiation and a direct effect of DHODH inhibition

Working on NB4 cells, a promyelocytic cell line, we observed for the first time that differentiation and apoptosis were partially independent effects of DHODH inhibition. Fig. 3A shows that, after 3 days of treatment, ATRA induced a strong expression of CD11c on leukemic cells (53.33% ± 2.02) while annexin expression was still minimal (10.4% ± 0.94), indicating that cells were differentiating but not dying yet. On the contrary, MEDS433 treatment greatly increased the percentage of apoptotic cells (43.43% ± 6.22), but it did not cause differentiation (1.97% ± 0.48). The combination of MEDS433 and ATRA induced differentiation (51.89% ± 10.53) and apoptosis (31.3% ± 3.79) simultaneously. Accordingly, the number of viable cells was modestly reduced when NB4 cells where just treated with ATRA, while it notably declined when cells were treated with MEDS433 alone or in combination with ATRA (Fig. 3A, right panel).

In all other tested cell lines, MEDS433 induced both apoptosis and differentiation; however, by analyzing the kinetics of these two phenomena, we noticed that the expression of annexin V anticipated the increase of the differentiation markers (Fig. 3B). These results indicate that at least a subset of cells undergoing apoptosis are not at the end of their differentiation process.

**Figure 3:**
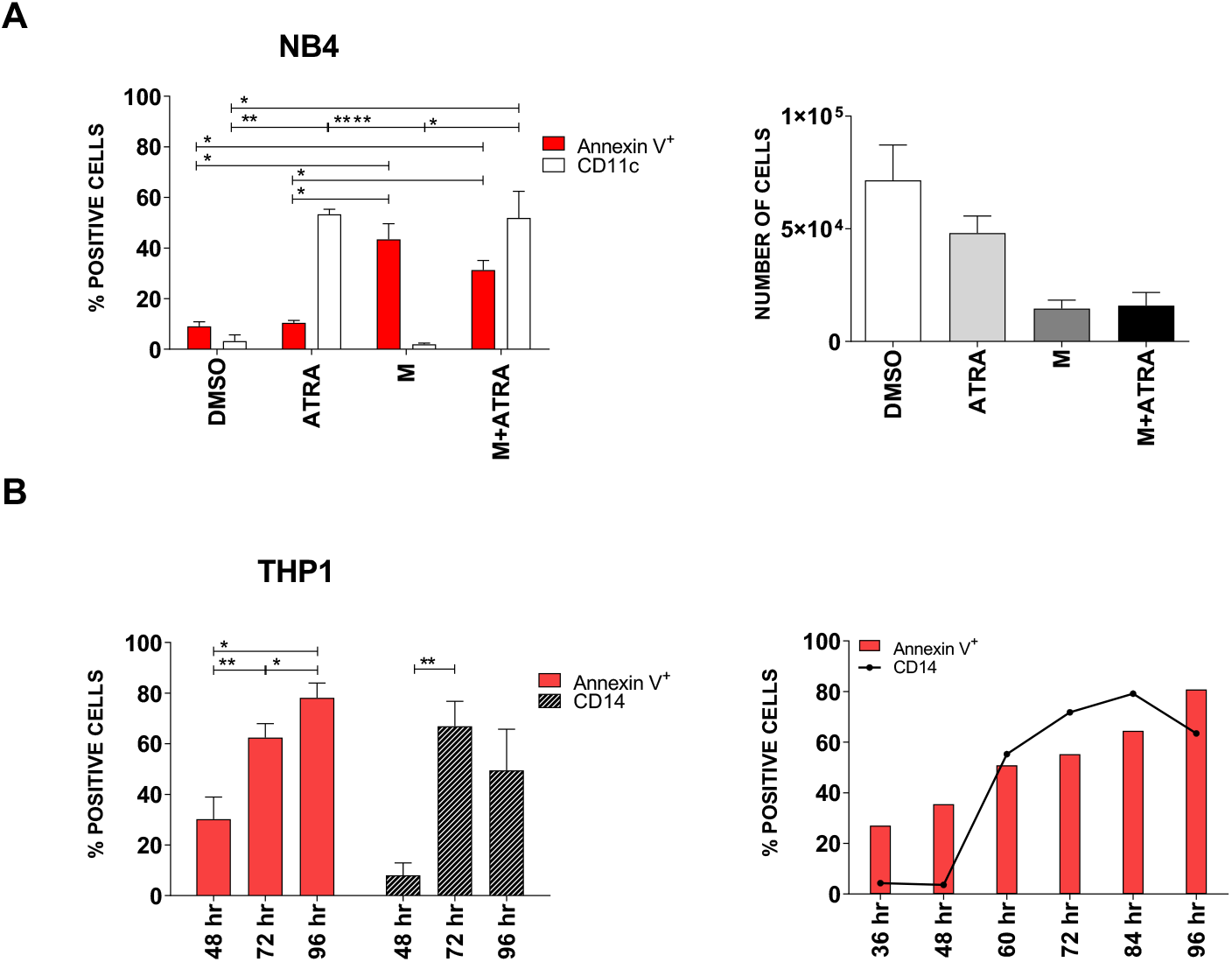
MEDS433 can induce apoptosis and differentiation separately. (A) Left panel: Apoptotic and differentiating rate of NB4 cells treated with ATRA, MEDS433 or both. Right panel: viable NB4 cells after treatment with ATRA, MEDS433 or both. (B) Kinetic of apoptosis and differentiation of THP1 cells treated with MEDS433 (one representative experiment out of 2 performed). In all the experiments, MEDS433 was utilized at 1 *μ*M and ATRA at 0.1 *μ*M. Differentiation of NB4 cells was best evaluated with CD11c expression, while THP1 cells with CD14 expression. The analyses shown in panel A were performed after 3 days of treatment. DMSO: dimethyl sulfoxide. ATRA: all-trans retinoic acid. M: MEDS433. hr: hours. (A) Statistical significance: Anova/Tukey, *p < 0.05; **p < 0.01; ***p < 0.001; ****p < 0.0001. (B) Statistical significance: t-test, *p < 0.05; **p < 0.01.

DHODH inhibitors, hence, work differently from other differentiating agents like ATRA, as they are able to directly cause apoptosis.

### 2.4 The combination of MEDS433 with classical antileukemic agents results in near-additive effects

In order to increase the apoptotic rate of AML cell lines, we first combined our DHODH inhibitor with chemotherapeutic agents that are classically used to treat AML, i.e., Ara-c, anthracyclines and decitabine. The concentrations of Ara-C (1 *μ*M), idarubicin (0.005 *μ*M) and decitabine (0.250 *μ*M) were chosen to induce a 15 to 40% apoptotic rate when administered alone, in order to explore synergistic combinations. Fig. 4 and Fig. S2 show that the combination of MEDS433 with Ara-C, idarubicin or decitabine significantly increases the apoptotic rate of either compound alone, resulting in a near-additive effect. However, in order to mimic *in vivo* conditions, we performed these experiments also in the presence of low, physiological, uridine levels. Fig. 4C shows that even low levels of uridine reduce the ability of MEDS433 to induce apoptosis, resulting in a substantial decrease of efficacy of MEDS433 alone and in combination with decitabine (from 69.32% ± 2.13 to 35.80% ± 1.13).

**Figure 4:**
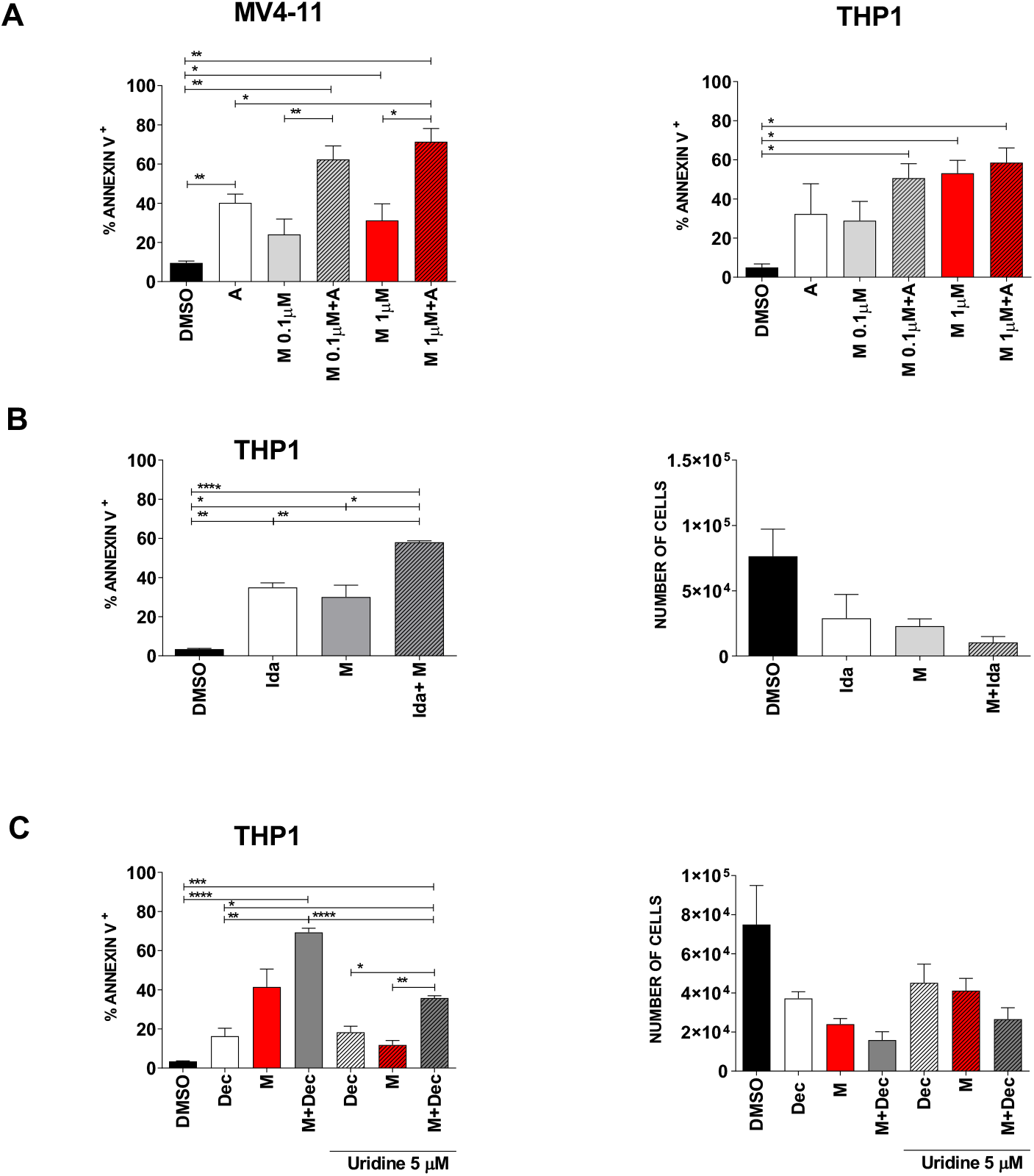
The combination of MEDS433 with classical antileukemic agents results in near-additive effects. (A) Apoptosis induced by MEDS433, Ara-C and their combination on MV4-11 (left panel) and THP1 (right panel) cells. (B) Apoptosis (left panel) induced by MEDS433, idarubicin and their combination on THP1 cells; the right panel represents the number of viable cells after the corresponding treatment. (C) Apoptotic rate (left panel) and cell viability (right panel) of THP1 cells treated with decitabine, MEDS433 or both, with or without physiological levels of uridine (5 *μ*M). If not otherwise specified, MEDS433 was utilized at 0.1 *μ*M. Apoptosis and cell counts were evaluated after 3 days of treatment. DMSO: dimethyl sulfoxide. M: MEDS433. A: Ara-C. Ida: idarubicin. Dec: decitabine. Statistical significance: Anova/Tukey, *p < 0.05; **p < 0.01; ***p < 0.001; ****p < 0.0001.

### 2.5 The combination of MEDS433 with hENT1/2 inhibitors results in synergistic effects against AML

Faced with results reported in Fig. 4C, we decided to investigate the antagonist role of uridine on DHODH inhibitors. Physiological uridine plasma levels are variable, approximately ranging from 2 to 6 *μ*M, being influenced by many factors, from physical activity to beer consumption [14, 15]. Fig. 5A shows that the apoptotic rate induced by MEDS433 was indeed reduced at increasing levels of uridine, potentially limiting the role of DHODH inhibitors in AML in vivo.

**Figure 5:**
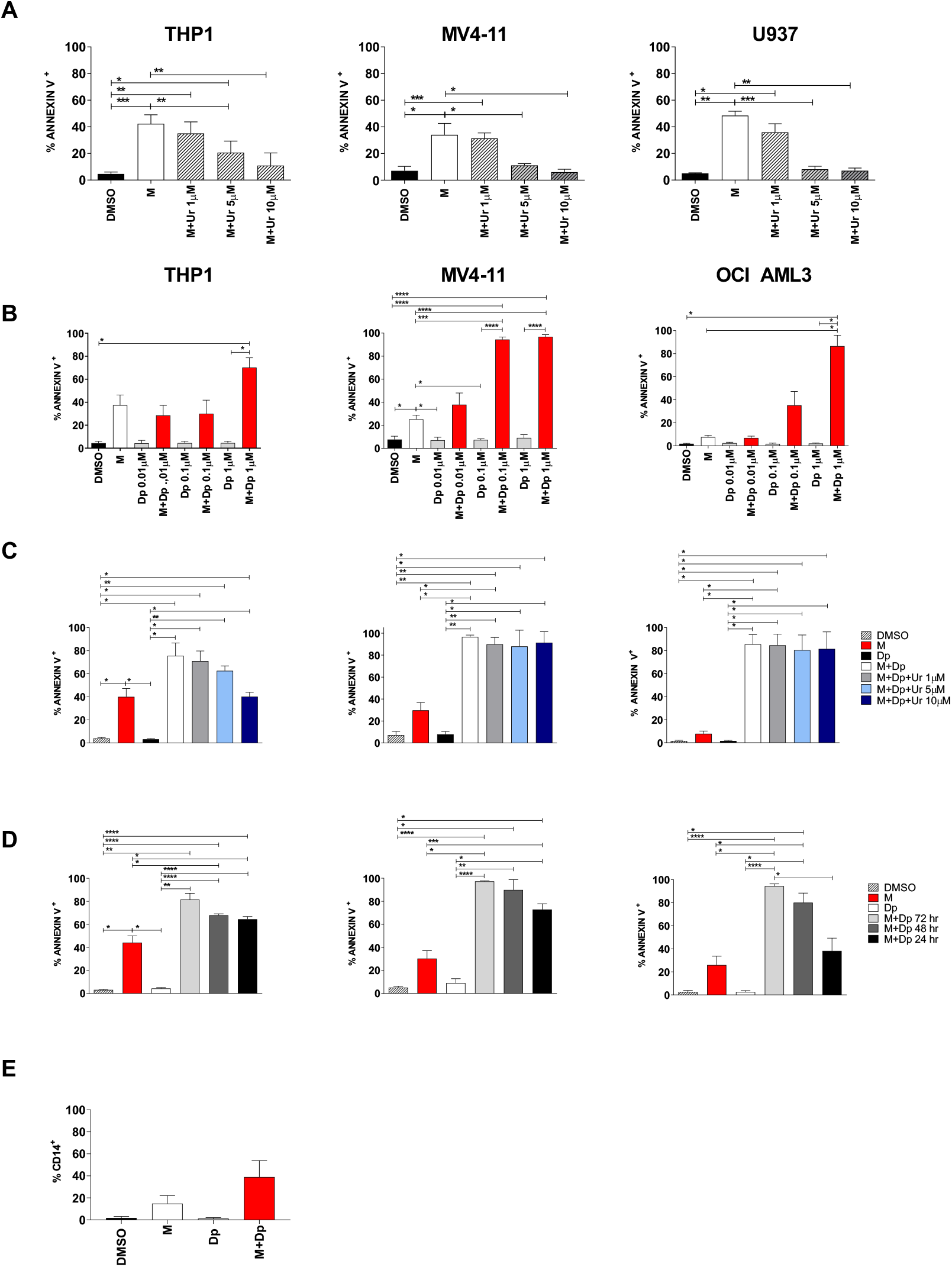
The combination of MEDS433 and dipyridamole results in synergistic effects. (A) Analysis of the apoptotic rate induced by MEDS433 on different cell lines, when utilized alone or in the presence of uridine at low concentrations (1 to 10 *μ*M). (B) Apoptosis induced by MEDS433, dipyridamole and their combination on THP1, MV4-11 and OCI-AML3 cells. (C) Apoptosis induced by MEDS433 in combination with dipyridamole in the presence of uridine at low concentrations (1 to 10 *μ*M) on THP1, MV4-11 and OCI-AML3 cells. (D) Cells were exposed to MEDS433 for 3 days and to dipyridamole for 3, 2 or 1 concomitant day(s), as shown in the figure legend; apoptosis was then evaluated on day 3. (E) Differentiation induced by MEDS433 and dipyridamole, alone or in combination, on THP1 cells; the differentiation analysis was performed on day 2. DMSO: dimethyl sulfoxide. M: MEDS433. Dp: dipyridamole. In all the experiments, MEDS433 was utilized at 0.1 *μ*M and apoptosis was evaluated after 3 days of treatment. For (B), (C), (D) and (E): THP1 in the left panel, MV4-11 in the middle panel, and OCI AML3 in the right panel. Statistical significance: Anova/Tukey, *p < 0.05; **p < 0.01; ***p < 0.001; ****p < 0.0001.

In order to minimize this issue, we decided to block both the de novo and the salvage pathway by combining MEDS433 to dipyridamole, a known inhibitor of nucleoside/nucleotide transport channels hENT1/2. As we were looking for a synergistic effect, we challenged our system using low concentrations of MEDS433 (0.1 *μ*M) and increasing concentrations of dipyridamole. Dipyridamole alone had no effect on any AML cell line, while MEDS433 confirmed its performances in the less favourable conditions (0.1 *μ*M). Notably, the combination of MEDS433 and dipyridamole resulted in a dramatic increase of the apoptotic rate, especially with dipyridamole concentrations above 0.1 *μ*M. All tested cell lines, even the most resistant to DHODH inhibitors, experienced this phenomenon (Fig. 5B and Fig. S3A). We also challenged this new combination by i) performing experiments in the presence of low uridine levels (Fig. 5C and Fig. S3B), and ii) adding dipyridamole for a limited amount of time (Fig. 5D): in both cases, the apoptotic rate was unaffected or largely preserved. Fig. 5E shows that the synergy of this combination was not limited to apoptosis, but rather extended to the differentiating effect as well. Unlike other experiments, the differentiation analysis had to be performed on day 2, resulting in worst results compared to Fig. 1D, because on day 3 the apoptotic rate was too high and compromised the reliability of results. Finally, in order to confirm the underlying mechanism, we substituted dipyridamole with dilazep, another hENT1/2 channel blocker. Fig. S3C shows that the combination of MEDS433 and dilazep has a strong apoptotic effect, similar to the dipyridamole and MEDS433 combination.

### 2.6 Uridine reversal: a matter of concentration and transporters

When uridine was added to the combination experiments (MEDS433 + dipyridamole) at hyperphysiological concentrations (100 *μ*M), apoptosis was almost totally abrogated in THP1, OCI AML3 and NB4 cells, and reduced in MV4-11 and U937 cells (Fig. 6A and Fig. S4); similar results were obtained when dipyridamole was substituted with dilazep (Fig. S3C). These data confirm that the combination induces apoptosis through pyrimidine starvation rather than off-target effects, but they also demonstrate that low dose dipyridamole is not able to fully block the salvage pathway. However, high dipyridamole concentrations (10 *μ*M) could restore almost totally the combination’s activity even with uridine 100 *μ*M, probably by inhibiting the vast majority of pyrimidine access channels (Fig. 6B). Similar results were obtained with dilazep (Fig. S3C). The reason why different cell lines have different sensitivity to the combination therapy, are differently rescued by high uridine concentrations and react differently to high dipyridamole concentrations is not clear; one hypothesis is that the sensitivity to pyrimidine depletion could inversely correlate with the levels of hENT2 protein, as suggested by expression data shown in Fig. 6C.

**Figure 6:**
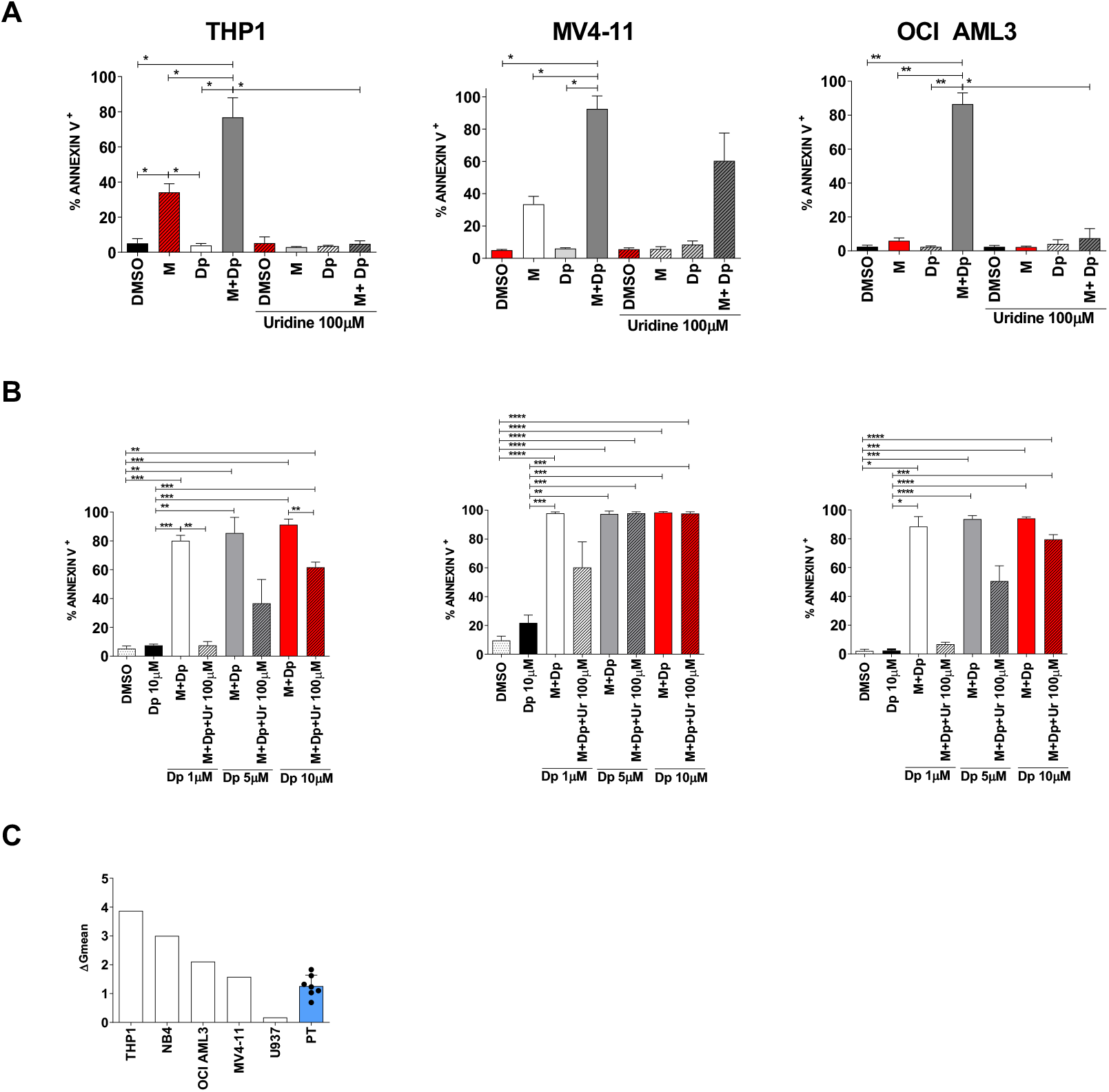
The uridine rescue is influenced by uridine concentrations and by the availability of active transporters. (A) Apoptosis induced by MEDS433, dipyridamole or their combination, with and without uridine at 100 *μ*M. Dipyridamole was utilized at the standard dose of 1 *μ*M. (B) Apoptosis induced by MEDS433 in combination with increasing dipyridamole concentrations, with and without uridine at 100 *μ*M. For (A) and (B): THP1 in the left panel, MV4-11 in the middle panel and OCI AML3 in the right panel. (C) hENT2 expression levels in utilized cell lines and primary AML cells, assessed by flow cytometry. DMSO: dimethyl sulfoxide. M: MEDS433. Dp: dipyridamole. Ur: uridine. PT: patients. In all the experiments, MEDS433 was utilized at 0.1 *μ*M and apoptosis was evaluated after 3 days of treatment. Statistical significance: Anova/Tukey, *p < 0.05; **p < 0.01; ***p < 0.001, ****p < 0.0001.

### 2.7 Approaching the clinical setting (I): effectiveness

Once disclosed the remarkable synergistic effect of MEDS433 with dipyridamole, we challenged this combination substituting our compound with teriflunomide, a 320-times less potent DHODH inhibitor, which is already commercialized as immunosuppressor. While teriflunomide alone did not have a remarkable apoptotic effect in any AML cell line, even at high doses, its combination with dipyridamole resulted in the net increase of the apoptotic rate far above 60%, in every tested AML cell line, even when *in vivo* conditions were mimicked (uridine 5 *μ*M) (Fig. 7A). Going further, we treated AML primary cells *in vitro:* again, while dipyridamole, MEDS433 or teriflunomide alone could not induce apoptosis, the combination of either DHODH inhibitor with dipyridamole resulted in a strong synergistic effect (Fig. 7B, Fig. 7C). In particular, of the 12 samples tested, 10 had an average apoptotic rate of 61.49% (range 43.92-82.40%) after just 3 days of treatment; the 2 remaining samples showed less impressive results (apoptotic rate of 18,3% and 26,3% respectively), demonstrating to be less sensitive but not completely non-responders. When this experiment was repeated in the presence of uridine 100 *μ*M, the apoptotic rate was largely reduced, excluding again off-target effects; however, high dipyridamole concentrations could almost totally rescue the apoptotic rate (Fig. 7B right panel). Noteworthy, all primary AML samples contained >75% blasts as detected by flow cytometry, and 10 out of 12 patients had a high-risk AML, whether for the European Leukemia Net risk score (FLT3 or Bcr/Abl mutated AML) or because they had therapy-related or secondary AML (post myelodysplastic or myeloproliferative syndromes), as shown in Tab. S2. On the whole, primary cells showed high sensitivity to the combination therapy, behaving similarly to OCI AML3 and MV4-11 cells. Not surprisingly, their hENT2 levels were comparable (Fig. 6C).

**Figure 7:**
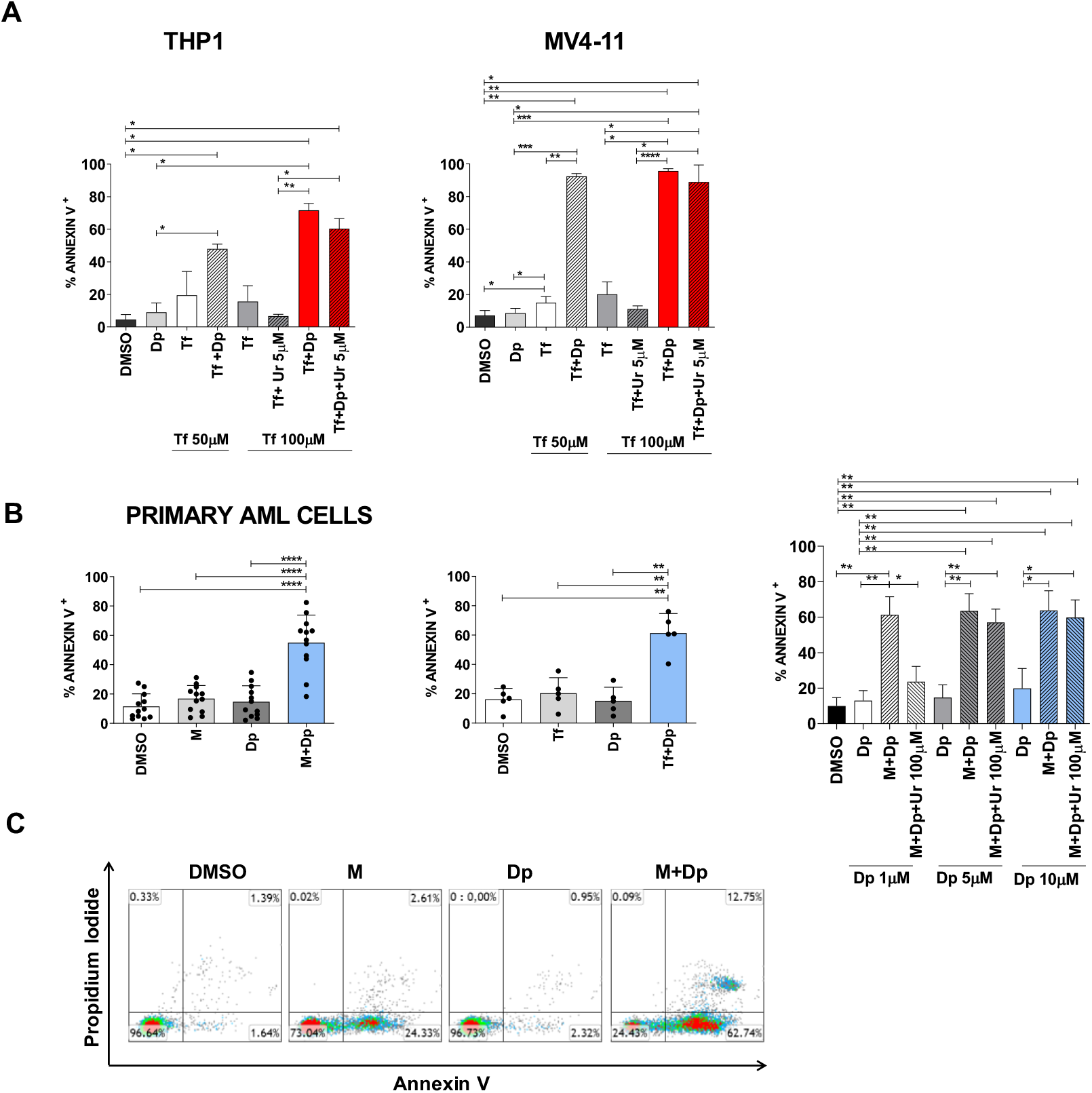
Dipyridamole synergizes with other DHODH inhibitors, and their combination is effective against primary AML cells. (A) Apoptosis induced by teriflunomide, dipyridamole and their combination, alone or in the presence of physiological uridine concentrations (5 *μ*M), on THP1 and MV4-11 cells. (B) Apoptosis induced by MEDS433 (left panel) or teriflunomide 100 *μ*M (middle panel), alone and in combination with dipyridamole, on AML primary cells. The right panel shows the apoptotic rate induced by MEDS433 alone and in combination with increasing dipyridamole concentrations, with or without uridine at hyperphysiological concentrations (100 *μ*M). (C) Flow cytometry plots on a representative sample of primary AML cells treated with MEDS433, dipyridamole and their combination (patient 3 of Tab. S2). In all the experiments, MEDS433 was utilized at 1 *μ*M and apoptosis was evaluated after 3 days of treatment; unless otherwise specified, dipyridamole was utilized at 1 *μ*M. DMSO: dimethyl sulfoxide. M: MEDS433. Dp: dipyridamole. Tf: teriflunomide. Ur: uridine. Statistical significance: Anova/Tukey, *p < 0.05; **p < 0.01; ***p < 0.001, ****p < 0.0001.

### 2.8 Approaching the clinical setting (II): toxicity

Blocking the *de novo* pyrimidine biosynthesis and/or the salvage pathway could be theoretically toxic even for non-cancer cells. Fig 8A shows that nor MEDS433 alone nor its combination with dipyridamole induced apoptosis on peripheral blood mononuclear cells (PBMC) or monocytes alone (Fig. 8A and Fig. S5A). Moreover, MEDS433 was not toxic on mesenchymal cells (Fig. 1D) and did not influence lymphocyte differentiation (Fig. 8A, middle panel). However, when lymphocytes were activated with phytohemagglutinin (PHA)/IL2/IL7, treatment with MEDS433 induced a significant increase of the apoptotic rate, which was enhanced by dipyridamole (Fig. 8B). This effect was confirmed when MEDS433 was substituted with teriflunomide, and was not attenuated in the presence of physiological uridine levels (from 75.26% ± 6.2 to 75.37% ± 5.86, Fig. 8B, middle and right panel). To compare the immunosuppressive potency of MEDS433+dipyridamole and chemotherapy, we exposed activated T-lymphocytes to a short course of antileukemic treatment (MEDS433+dipyridamole vs Ara-C vs idarubicin), and then observed their apoptotic rate right after the exposure (day 3) and after 6 days of recovery in the presence of uridine at physiological levels. Fig. 8C shows that the combination of MEDS433+ dipyridamole has a strong immunosuppressive effect, comparable to chemotherapeutic agents, but far more sustained over time.

**Figure 8:**
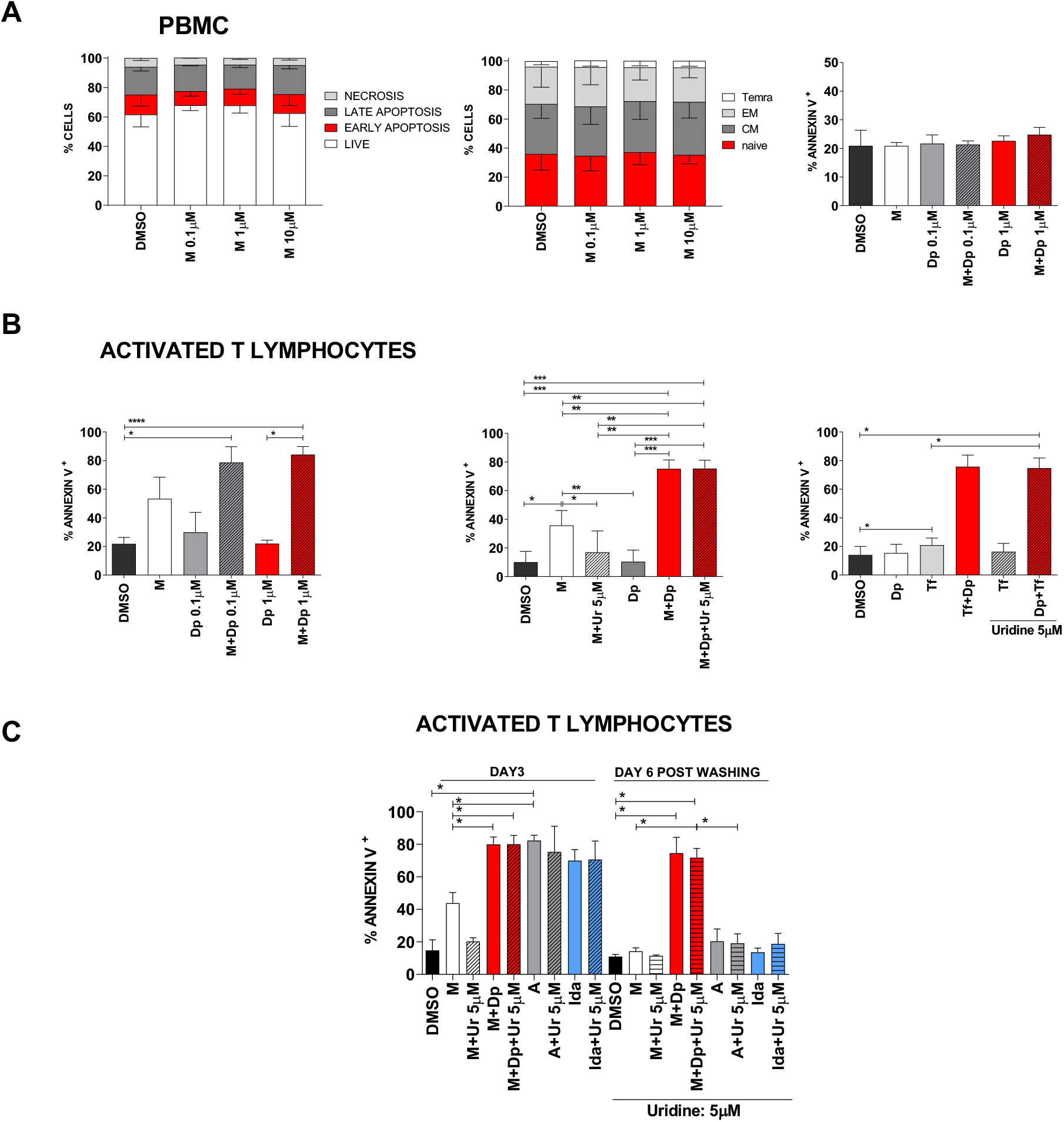
Toxicity of DHODH inhibitors alone and in combination with dipyridamole. (A) Left panel: apoptosis distribution induced by MEDS433 on PBMC, based on annexin V vs propidium iodide expression. Middle panel: differentiation of T-lymphocytes treated with MEDS433 at increasing concentrations; the analysis was performed on the CD3+ population, identifying the following subsets: naïve (CD45RA+CD62L+), CM (central memory: CD45RA-CD62L+), EM (effector memory: CD45RA-CD62L-), and Temra (terminally differentiated effector memory CD45RA+CD62L-). Right panel: apoptosis induced by MEDS433 at 0.1 *μ*M, dipyridamole and their combination on PBMC. (B) Apoptosis induced by MEDS433 alone (0.1 *μ*M) or in combination with dipyridamole on activated T-lymphocytes, without (left panel) or in the presence of uridine at the physiological concentration of 5 *μ*M (middle panel). The same experiment was performed by replacing MEDS433 with Teriflunomide 100 *μ*M (right panel). (C) Comparison between the long and short-term toxicity of MEDS433 (0.1 *μ*M) vs chemotherapy (Idarubicin or Ara-c) on activated T-lymphocytes. The cells were treated for 3 days with MEDS433 alone and in combination with dipyridamole, or with Ara-C or idarubicin. After 3 days, cells were washed and replated without drugs, in the presence of physiological uridine concentrations (5 *μ*M). If not otherwise specified, apoptosis was evaluated after 3 days of treatment. PBMC: peripheral blood mononuclear cells. DMSO: dimethyl sulfoxide. M: MEDS433. Dp: dipyridamole. Tf: teriflunomide. Ur: uridine. A: Ara-C. Ida: idarubicin. Statistical significance: Anova/Tukey, *p < 0.05; **p < 0.01; ***p < 0.001, ****p < 0.0001.

## 3 Discussion

The major advances of the last years in understanding AML biology have been essential to develop new targeted drugs, such as FLT3 and IDH1/2 inhibitors. However, these “mutation-specific” drugs only address one clone, while we have learnt that AML is often an extremely heterogeneous and dynamic disease, with multiple subclones emerging or disappearing depending on their selective advantage. Therapeutic agents are a major variable influencing this selective advantage: mutation-specific drugs, if used alone, generally induce clonal escape; chemotherapeutic drugs are extremely effective on proliferating cells, but quiescent leukemic stem cells could be spared, leading to disease relapse; moreover, chemo-resistant clones, e.g. those harbouring p53 mutation or complex karyotypes, are often selected.

For this reason, the wise combination of multiple synergic drugs or approaches could be fundamental to efficiently target AML and avoid the emergence of resistances. Differentiating and pro-apoptotic drugs could be essential ingredients of these combinations as they could i) force immature cells to differentiate, increasing their exposure to other drugs, being potentially active on leukemic stem cells, and ii) be effective on a wide variety of subclones, not being mutation specific. Not surprisingly venetoclax, a bcl-2 inhibitor with strong pro-apoptotic effect, is an emerging and extremely promising drug in the field, especially when combined with other drugs [16].

In our work, we decided to focus on another non-mutation-restricted mechanism: the metabolism. It has long been known that oncogenes deviate several metabolic pathways to support cancer growth, leading to metabolic addiction [17]; vice versa, the dysregulation of metabolism can be directly tumorigenic [18]. Since the first attempts in the 1940s [19], classical antimetabolites (anti-folate drugs, pyrimidine or purine analogues) have represented a mainstay of cancer therapy.

In 2016, some compounds that induce pyrimidine depletion, i.e., DHODH inhibitors, were found to promote differentiation in several AML models [1], underlying the wide influence of metabolism on cancer cells behaviour. From that moment, the field rapidly expanded, with multiple research teams creating new agents that joined brequinar, an old but potent DHODH inhibitor, in clinical trials. Our group has recently developed MEDS433, a DHODH inhibitor which is able to induce differentiation at a 1-log lower concentration compared to brequinar [2]. Given its mechanism of action, and previous results in solid tumors, we hypothesized that DHODH inhibition could directly influence apoptosis, so we started to investigate this phenomenon in AML. Basically, all tested AML cell lines underwent apoptosis when treated with MEDS433, with the less sensitive cell lines responding to a prolonged exposure. Similarly to the differentiation setting, MEDS433 could induce apoptosis at a 1-log lower concentration compared to brequinar, and apoptosis was totally abrogated when uridine was added at hyperphysiological concentration, indicating that apoptosis is a specific, on-target effect. Apoptosis was preserved in niche-like conditions, i.e., when cell lines were co-cultured with stromal cells and in hypoxic conditions: this finding is in line with data shown by Sykes et al, which observed a decrease in the number of self-renewing cells *in vivo* [1]. Finally, analysing the kinetics of differentiation and apoptosis as well as the effect of DHODH inhibition on a promyelocytic cell line, we demonstrated that apoptosis is not a mere consequence of differentiation but, on the contrary, it is both the result of differentiation and a direct effect of DHODH inhibition.

Being able to induce both differentiation and apoptosis, and targeting a non-mutation-restricted metabolic pathway, DHODH inhibitors could represent the perfect drugs to obtain synergic treatments in AML, so we started to investigate new combinations. Indeed, several studies in solid tumors have previously shown that DHODH inhibition could sensitize cancer cells to chemotherapy, eventually overcoming chemoresistance [20, 21, 22].

As AML is conventionally treated with an anthracycline (daunorubicin or idarubicin) and a pyrimidine analogue (Ara-c), we combined MEDS433 with these drugs. In both cases, the apoptotic rate was significantly increased, suggesting that the addition of DHODH inhibitors to classical chemotherapy could improve the performances of the traditional 3+7 induction regimen for AML, and maybe overcome chemoresistance. Indeed, Imanishi et al have already shown that DHODH inhibition could restore sensitivity to 5-azacytidine in resistant cell lines, *in vitro* [23].

While these combinations yielded promising results, our major concern was the *in vivo*, and most importantly, the “in human” performance of DHODH inhibitors, due to the physiological presence of uridine in patients’ plasma and the possibility for cancer cells to rely on the salvage pathway for pyrimidine supply. An incomplete pyrimidine starvation, in fact, is thought to be one of the major causes of DHODH inhibitors failure in clinical trials [10, 11, 12]. Indeed, we demonstrated that MEDS433 showed less impressive results when uridine was added at physiological levels. In order to block both the *de novo* and the salvage pathway, we combined MEDS433 to dipyridamole, a known inhibitor of nucleoside/nucleotide transport channels.

This association resulted in the dramatic increase of the apoptotic rate in all tested cell lines; besides, the synergy applied also to the differentiation process, making this combined treatment a reminiscent of ATRA and arsenic therapy in promyelocytic leukemia.

The block of both pyrimidine sources retained its effectiveness even when MEDS433 was substituted with a much less potent DHODH inhibitor, teriflunomide, which is already commercialized for multiple sclerosis. More importantly, the combination of MEDS433 with dipyridamole, but not the individual compounds, was extremely effective in inducing apoptosis in several AML primary cells: 10 out of the 12 tested samples were highly sensitive, with a median apoptotic rate exceeding 60% after just 3 days of treatment, despite being characterized for the vast majority by a high-risk profile.

Preliminary data suggest that the sensitivity to our combination therapy could correlate with the abundance of hENT2 channels, i.e., with the ability of the cells to salvage pyrimidines. Noteworthy, all primary AML cells tested had intermediate-low hENT2 levels, endorsing this hypothesis.

Although our results will need confirmatory experiments, we can already predict, based on our *in vitro* data, that the effect of the combined treatment on AML cells should not be affected by physiological uridine levels *in vivo*. Indeed, the pro-apoptotic effect of MEDS433 + dipyridamole on AML cell lines was unchanged or negligibly reduced when experiments were performed at physiological levels of uridine. More importantly, on primary AML cells, we demonstrated that the combination of MEDS433 with slightly higher dipyridamole concentrations (between 1 and 5 *μ*M) could still induce apoptosis in extreme conditions, i.e, with uridine concentrations 20-50 times more than normal. This large margin of safety represents a guarantee against possible escape mechanisms that leukemic cells could play out, such as transporter upregulation.

Given this dramatic effectiveness on several AML cells, we investigated whether this combination could be toxic also for non-cancer cells. Our data show that toxicity is limited to proliferating cells, and in particular activated T-lymphocytes, which is consistent with the use of DHODH inhibitors as immunosuppressors. While this effect would be undesirable during induction/consolidation treatments, it could be extremely useful in the post-transplant setting. Drugs targeting graft-versus-host disease (GVHD) are generally immunosuppressive and potentially favour relapse by inhibiting graft-versus-leukemia; conversely, a suboptimal control of alloreactive T-lymphocytes could repress leukemia but trigger severe GVHD. Our combination therapy, on the contrary, could target leukemia while suppressing activated lymphocytes.

In the 1990s, a limited number of studies investigated the combination of drugs inhibiting both the salvage pathway and the *de novo* synthesis in solid tumors: they obtained quite promising results *in vitro* but limited effectiveness in clinical trials [24, 25]. Several hypotheses were formulated to explain this failure [26], including an incomplete inhibition of the pyrimidine supply, whether for a wrong brequinar administration schedule or because other channels allowed minimal quantity of uridine to enter the cells. Another hypothesis is that this combination was cytostatic in cancer cells. Our data show that this combination is highly effective in AML, where it has cytotoxic effects. As a matter of fact, the kinetics of solid and hematological tumors, and especially AML, are extremely different; moreover, the pyrimidine starvation induces differentiation in the AML settings, but not in solid tumors. Taken together these results suggest that the combination therapy of DHODH inhibitors and hENT blockers could work in AML much better than in solid tumors.

Finally, it is possible to hypothesize, in the future, a triple combination of chemotherapy, DHODH inhibitors and dipyridamole. In this context, finding the correct schedule of administration will be fundamental to maximize the synergism and the pyrimidine depletion, while minimizing toxicity. A triple combination, especially if given continuously, could be toxic even for non-cancer cells; moreover, the block of nucleotide transporters could prevent some chemotherapeutic drugs to enter the cells, such as decitabine or Ara-C. However, a wise schedule of administration could further enhance the antileukemic activity and overcome chemo-resistance. Just to mention few examples, a pre-treatment with DHODH inhibitors and dipyridamole could *i*) reduce the cytidine triphosphate (CTP) pool, sensitizing leukemic cells to azacytidine and decitabine by decreasing the competition for their incorporation into nucleic acids; and ii) force leukemic cells to increase the expression of hENT channels, potentially promoting the access of decitabine, azacytidine or Ara-C into the cells.

## 4 Conclusions

AML is an aggressive disease which is still characterized by a dismal prognosis in too many patients. Its biology reveals an heterogenous background, with multiple clones competing for selective advantage, but with some common features: a differentiation block, a reduced apoptosis and a metabolic addiction.

In this work we present a new combination, based on deep pyrimidine starvation and characterized by differentiating and pro-apoptotic features, that was effective on all AML cell lines and the vast majority of primary AML cells, with very different genetic backgrounds.

At the best of our knowledge, this is the first study investigating this combination treatment in AML; more importantly, as it addresses a completely new pathway, this treatment could be furtherly associated with classical antileukemic drugs, in order to find the best combination to treat AML.

## 5 Materials and Methods

### 5.1 Reagents

MEDS433 and Brequinar were synthesized as described in [2]. ATRA, Teriflunomide, Dilazep and uridine were purchased from Sigma-Aldrich. Reagents were dissolved in DMSO and diluted in culture medium before use. Final DMSO concentration did not exceed 0.1%. Dilazep was dissolved in water. Ara-C (Citarabina Hospira, IL, Usa), Idarubicin (Zavedos, Pfizer, NY, Usa), Dipyridamole (Persantin, Boehringer Ingelheim, Germany), Decitabine (Dacogen, Janssen) were purchased.

### 5.2 Cell culture

All cell lines were purchased by DSMZ. The human cell lines THP1 (acute monocytic leukemia M5), U937 (pro-monocytic myeloid leukemia), NB4 (promyelocytic leukemia M3), MV4-11 (acute monocytic leukemia M5) and OCI AML3 (acute myelomonocytic leukemia M4) were maintained in RPMI 1640 (Gibco, Thermo Fisher Scientific) supplemented with 10% heat-inactivated fetal bovine serum (FBS, Gibco, Thermo Fisher Scientific). The human stromal cell line HS-5 was cultured in DMEM (Microtech, Naples, Italy) supplemented with 10% FBS. Media were supplemented with 1% penicillin/streptomycin (Gibco, Thermo Fisher Scientific) and all cultures were maintained at 37°C, 5% CO_2_. Cell lines were maintained in culture for no longer than 5-6 weeks and were routinely tested for mycoplasma contamination. The genetic alterations of cell lines are shown in Tab. S1. The doubling time of the cell lines was available in the DSMZ catalogue

All cells lines were plated into 96-well round bottom plates at 1 × 10^4^ per well and treated with DMSO or a DHODH inhibitor: MEDS433 from 0.1 *μ*M to 10 *μ*M, Brequinar from 0.1 *μ*M to 10 *μ*M or Teriflunomide from 50 *μ*M to 100 *μ*M for three days. Some experiments were repeated in the presence of uridine (concentration range: 1 *μ*M to 100 *μ*M). For drug combinations, cell lines were treated with ATRA 0.1 *μ*M or Ara-C 1 *μ*M or Idarubicin 0.005 *μ*M or Decitabine 0.250 *μ*M or Dypiridamole (0.01 *μ*M-10 *μ*M). All experiments were repeated at least three times.

For hypoxia, 1 × 10^4^ THP1 cells were seeded into 96-well round bottom plates in a volume of 200 *μ*l of complete RPMI. Cell cultures were grown under hypoxic conditions using a Hypoxia Chamber (STEMCELL Technologies, Vancouver, Canada) at 37°C, 5% CO_2_ and 1% O_2_. MEDS433 was added after 24 hours adaptation period to the hypoxic conditions. Cells were treated under hypoxic conditions for three days. Analysis of viability and differentiation were performed at the end of cultures.

### 5.3 Co-culture

Stromal cell monolayers were generated seeding 1 × 10^3^ HS-5 in 96-well flat bottom tissue culture plates using IMDM (Microtech, Naples, Italy), supplemented with 10% FBS. A total of 1 × 10^4^ THP1 were seeded on the stromal cell monolayer after 24 hours and subsequently treated with MEDS 443 (1 *μ*M). After three days, the viability of THP1 was evaluated by Annexin V/Propidium Iodide (PI) assay (Miltenyi Biotec, Italy). Cells in suspension were collected and pooled with adherent cells treated with Trypsin/EDTA (Sigma Aldrich, Milan, Italy). Cells were incubated with AnnexinV-FITC and CD4 APC (Miltenyi Biotec, Italy) for 15 minutes in the dark at room temperature. After PI addition, cells were acquired on FACSVerse (BD-Biosciences San Jose-CA) and THP1 cells were isolated from stroma cells by expression of CD4.

### 5.4 Primary cells

Primary AML cells were obtained from peripheral blood or bone marrow of patients hospitalized in the University Division of Hematology and Cell Therapy of A.O. Ordine Mauriziano, Turin, Italy, after informed consent. Samples collection and analyses were approved by the Local Ethical Committee (Deliberation number 602). All samples contained >75% blasts as detected by flow cytometry and the characteristics of patients are shown in Tab. S2. Fresh or frozen samples were used. Mononuclear cells were purified by Ficoll-Hypaque density gradient centrifugation (Sigma Aldrich), and PBMC were cultured in IMDM (Microtech, Naples Italy) supplemented with 20% FBS, insulin-transferrin-selenium (Gibco, Thermo Fisher Scientific), FLT3L (20 ng/ml Miltenyi Biotec), SCF (20 ng/ml Miltenyi Biotec), and G-CSF (100 ng/ml, Nivestim, Hospira IL, Usa).

Mononuclear cells were isolated by Ficoll-Hypaque gradient from healthy donor after informed consent. PBMC were maintained in complete RPMI 1640 and treated directly with MEDS433 alone or with Dypiridamole (0.1–1 *μ*M) or stimulated with 2% phytohemagglutinin (PHA, Gibco, Thermo Fisher Scientific), IL2 200U/ml (Peprotech, Inc), and IL7 10ng/ml (Miltenyi Biotec) for 72 hours. After stimulation, cells were washed and re-suspended in complete RPMI 1640 supplemented with fresh cytokines and exposed to MEDS433 (0.1 *μ*M), Idarubicin (0.005 *μ*M), Ara-C (1 *μ*M) for three days. Analysis of viability was performed by AnnexinV assay. To evaluate the lasting effect of the drugs on viability, cells were washed and re-plated in medium supplemented with fresh cytokines and uridine 5 *μ*M and cultured for additional 6 days.

### 5.5 Apoptotic assay

Apoptosis was examined by flow cytometry using Annexin V-FITC Kit (Miltenyi Biotec, Italy), according to the manufacturer’s instructions. Alternatively, cell lines or primary cells were incubated with Annexin V-FITC and CD14 APC (BD Biosciences) or CD11c APC (Thermo Fisher Scientific) for 15 minutes at room temperature in the dark. After the PI incorporation, samples were acquired on FACSVerse and analysed by Kaluza software version 1.2 (Beckman Coulter Fullerton, CA).

### 5.6 Flow Cytometry

PBMC samples were seeded (1 × 10^4^ cells/well) in complete medium with or without the indicated concentrations of compounds. After 72 hours incubation, cells were washed with PBS and stained with CD45RA PE (BD Biosciences), CD62L APC (Miltenyi Biotec), CD3 FITC (Miltenyi Biotec) at room temperature for 20 minutes. Data collection was done on FACSVerse and dead cells were excluded from the analysis, based on the use of PI (Sigma-Aldrich, Milan, Italy).

Cells lines or AML primary cells were incubated with a SLC29A2 polyclonal antibody (Thermo fisher Scientific) using a dilution of 1:25 for 20 minutes at room temperature after fixation and permeabilization. The secondary antibody (Cell signaling Technology USA, Danvers, MA) was used with a dilution of 1:200 for 20 min. All data were processed with Kaluza software version 1.2.

### 5.7 Trypan blue staining assay

For each experiment, all cells lines (1 × 10^4^) were treated for 3 days to the indicated concentrations of drugs in a final volume of 200 *μ*l. Control groups were maintained in the same culture condition with DMSO. The total number of viable cells was determined by trypan blue staining (Sigma-Aldrich).

### 5.8 Statistical analysis

Statistical analysis was either performed using Prism software, version 5.0 (GraphPad Software, San Diego, CA), as well as a Python script that relied on the open-source SciKit Learn library. All data is reported as means ± SD with at least three independent experiments. Two tail paired Student’s t-test was calculated to assess differences between two groups and p < 0.05 was considered significant. For multiple comparisons, one-way ANOVA tests were performed and combined with Tukey’s tests for *post hoc* analyses. Also in this case, a p-value < 0.05 was considered significant. Due to the high complexity of the figures, in some panels the statistical significance is shown only for the most relevant biological comparisons.

The linear regression that correlates the doubling time of the cells lines with the apoptotic rate was performed through the SciKit Learn library, and the obtained segment represents a linear interpolation between the means of the observations.

## Abbreviations

DHODH: Dihydroorotate Dehydrogenase
AML: Acute Myeloid Leukemia
ATRA: All-Trans Retinoic Acid
hENT1/2: human Equilibrative Nucleoside Transporters
hCNT: human Concentrative Nucleoside Transporters
DMSO: Dimethyl Sulfoxide
PI: Propidium Iodide
M: MEDS433
BRQ: Brequinar
Ur: Uridine
A: Ara-C
Ida: Idarubicin
Dec: Decitabine
Dp: Dipyridamole
Dz: Dilazep
Tf: Teriflunomide
GVHD: Graft-Versus-Host Disease
CTP: Cytidine Triphosphate
FBS: Fetal Bovine Serum
PHA: Phytohemagglutinin

## Availability of data and materials

Experimental data are available from the authors upon reasonable request and with permission of the University of Turin.

## Ethics approval and consent to participate

Samples collection and analyses were approved by the Ethical Committee of the A.O. Ordine Mauriziano (Deliberation number 602). All patients gave their informed consent for the use of their biological material and for publications of the obtained results.

## Authors’ contributions

V.G., A.C., G.S and P.C conceptualized the project; V.G. and P.C. wrote the manuscript; M.H., N.V., G.C, A.M., S.R. and P.C performed the biological experiments; V.G., A.C., G.S, M.H., N.V., S.R., G.C., A.M. and P.C. designed the experiments and discussed the biological results; V.T. and P.C performed statistical analyses; S.S., A.C.P., M.G., D.B. an M.L.L. designed and developed MEDS433. All co-authors contributed to the final version of the manuscript.

