## Supplementary Figures and Tables for "The Synergism Between DHODH Inhibitors and Dipyridamole Leads to Metabolic Lethality in Acute Myeloid Leukemia"

(Dated: October 13, 2020)

### SUPPLEMENTARY FIGURES

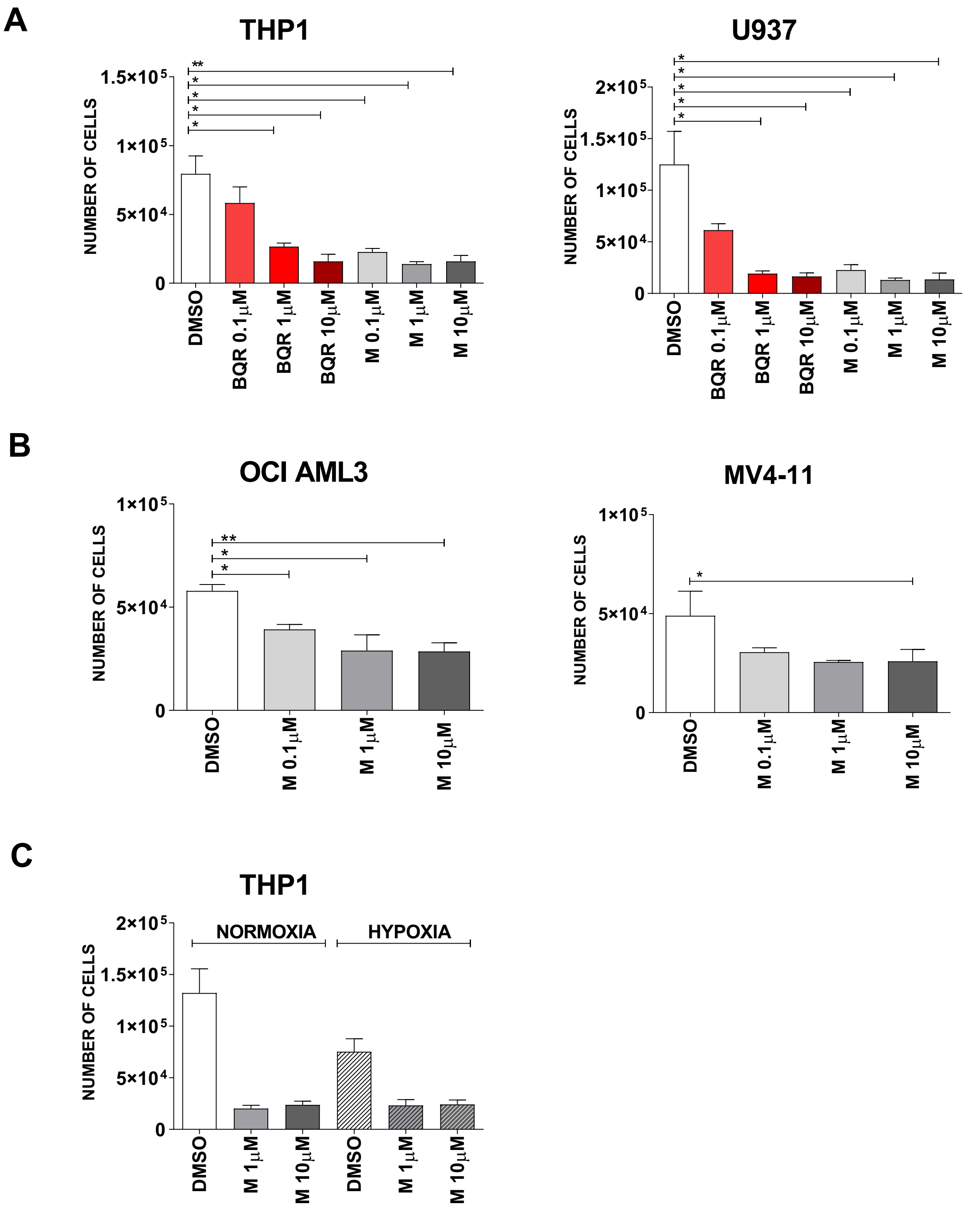


FIG. S1: (A) Comparison between the number of viable THP1 and U937 cells treated with MEDS433 or brequinar. (B) Reduction of OCI AML3 and MV4-11 cell viability after treatment with MEDS433 at increasing concentrations. (C) Evaluation of viable THP1 cells treated with MEDS433 in normoxic and hypoxic conditions. The analyses were performed after 3 days of treatment. DMSO: dimethyl sulfoxide. M: MEDS433; BQR: brequinar. Statistical significance: t-test, *$p < 0.05$; **$p < 0.01$; ***$p < 0.001$.

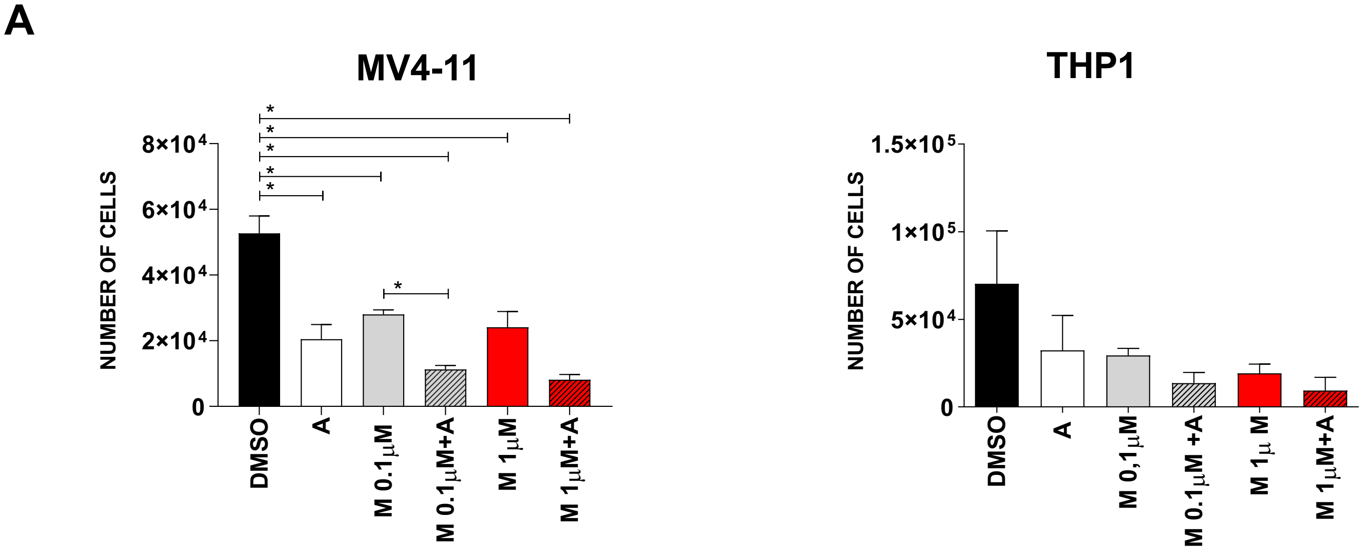


FIG. S2: (related to Fig. 4). MV4-11 (left panel) and THP1 (right panel) cells viability after treatment with MEDS433, Ara-C and their combination. The analyses were performed after 3 days of treatment. DMSO: dimethyl sulfoxide. M: MEDS433. A: Ara-C. Statistical significance: Anova/Tukey, *$p < 0.05$; **$p < 0.01$; ***$p < 0.001$; ****$p < 0.0001$.

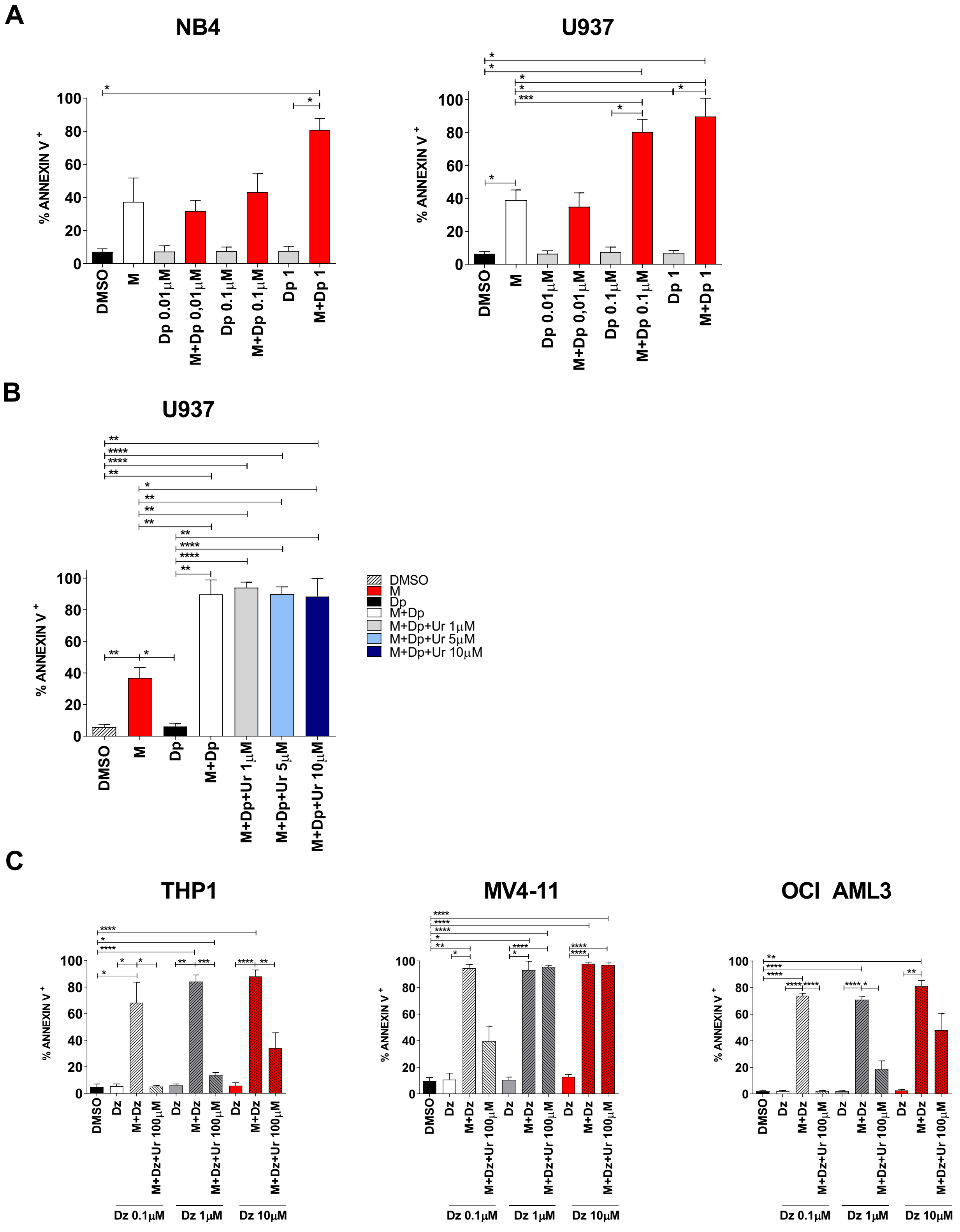


FIG. S3: (related to Fig. 5). (A) Apoptosis induced by MEDS433, dipyridamole and their combination on NB4 and U937 cells. (B) Apoptosis induced by MEDS433 in combination with dipyridamole in the presence of uridine at low concentrations (1 to 10 $\mu$M) on U937 cells. (C) Apoptosis induced by MEDS433, dilazep or their combination, with and without uridine at 100 $\mu$M, on THP1, MV4-11 and OCI AML3 cells. Dilazep was utilized at increasing concentrations, as shown in the figure. DMSO: dimethyl sulfoxide. M: MEDS433. Dz: dilazep. Ur: uridine. Apoptosis was evaluated after 3 days of treatment. Statistical significance: Anova/Tukey, *p $<$ 0.05; **p $<$ 0.01; ***p $<$ 0.001; ****p $<$ 0.0001.

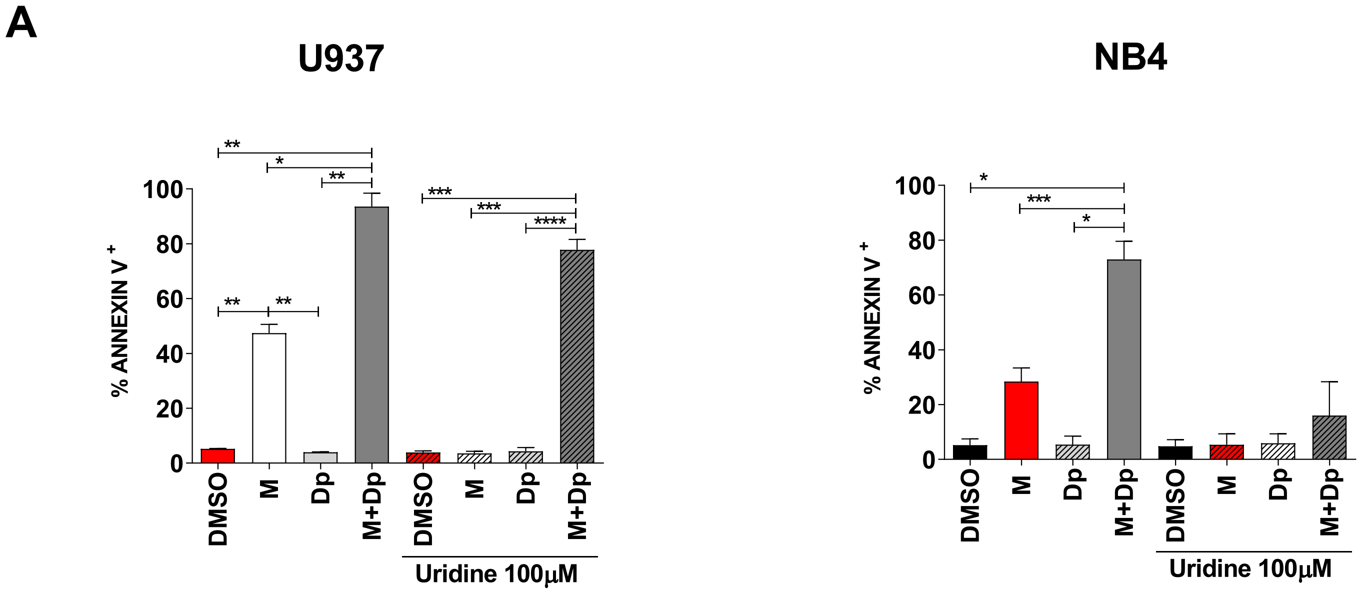


FIG. S4: (related to Fig. 6) Apoptosis induced by MEDS433, dipyridamole or their combination, with and without uridine at 100 $\mu$M, on U937 and NB4 cells. DMSO: dimethyl sulfoxide. M: MEDS433. Dp: dipyridamole. Ur: uridine. In all the experiments, MEDS433 was utilized at 0.1 $\mu$M, dipyridamole at 1 $\mu$M and apoptosis was evaluated after 3 days of treatment. Statistical significance: Anova/Tukey, *p < 0.05; **p < 0.01; ***p < 0.001; ****p < 0.0001.

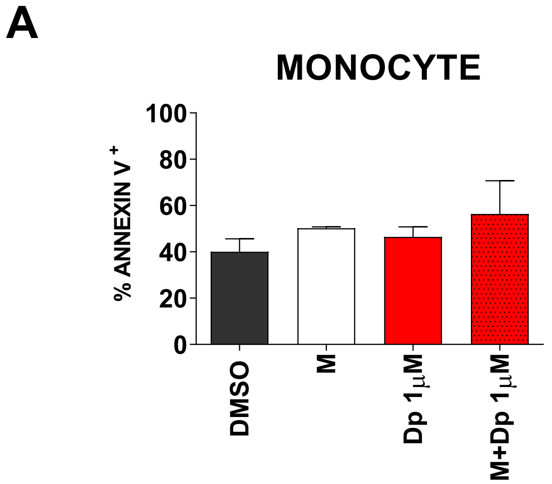


FIG. S5: (related to Fig. 8). Apoptosis of monocytes treated with MEDS433 alone (0.1 $\mu$M) or in combination with dipyridamole (1 $\mu$M) . Apoptosis was evaluated after 3 days of treatment. DMSO: dimethyl sulfoxide. M: MEDS433. Dp: dipyridamole. Statistical significance: Anova/Tukey, *p < 0.05; **p < 0.01; ***p < 0.001; ****p < 0.0001.

### SUPPLEMENTARY TABLES

| Cell line | Genetic alterations |
|---|---|
| U937 | CALM-AF10, PTEN, TP53 |
| THP1 | NRAS, TP53, MLL-AF9, CSNK2A1-DDX39B |
| OCI AML3 | DNMT3A, NPM |
| NB4 | PML-RARa, KRAS, TP53 |
| MV4-11 | MLL-AFF1, FLT3-ITD |

TABLE S1: Molecular alterations of utilised cell lines. All cell lines are characterized by complex karyotypes.

| PT | TYPE OF SAMPLE | WHO classification - genetic alterations |
|---|---|---|
| 1 | PB | AML with myelodysplasia-related changes |
| 2 | PB | AML with myelodysplasia-related changes |
| 3 | BM | Therapy-related myeloid neoplasms |
| 4 | BM | AML NOS, without maturation |
| 5 | PB | AML NOS, Acute myelomonocytic leukemia |
| 6 | BM | AML NOS, Acute myelomonocytic leukemia (FLT3+) |
| 7 | PB | AML NOS, without maturation (secondary to MPN, myeloproliferative neoplasm) |
| 8 | PB | Provisional entity: AML with BCR-ABL1 (blast crisis of CML) |
| 9 | PB | AML NOS, Acute myelomonocytic leukemia (FLT3+) |
| 11 | PB | AML NOS, without maturation (secondary to MPN, myeloproliferative neoplasm) |
| 10 | PB | AML NOS, Acute myelomonocytic leukemia (FLT3+) |
| 12 | PB | AML NOS, Acute myelomonocytic leukemia (FLT3+) |

TABLE S2: WHO classification and molecular alterations of utilized AML samples.
